# Naturalistic Stimulus Reconstruction from fMRI: A Primer in the Natural Scenes Dataset

**DOI:** 10.64898/2026.03.26.714100

**Authors:** Umur Yıldız, Burcu A. Urgen

**Affiliations:** Department of Neuroscience, Bilkent University, Bilkent, Ankara, Türkiye; Department of Psychology, Bilkent University, Bilkent, Ankara, Türkiye; Aysel Sabuncu Brain Research Center, Bilkent University, Ankara, Türkiye

**Keywords:** fMRI, decoding, reconstruction, NSD, CLIP, diffusion

## Abstract

Reconstructing natural images from brain activity represents one of the most compelling demonstrations of the synergy between modern neuroimaging and machine learning. However, the computational pipelines underlying these results remain scarcely accessible, difficult to reproduce, and offer limited opportunities for hands-on experimentation. They depend on large codebases, expensive hardware, and multiple representational stages whose interactions are not obvious. We present a step-by-step tutorial, organized across six notebooks, for reconstructing natural images from fMRI responses in the Natural Scenes Dataset. The workflow walks the reader through three main stages: predicting coarse image structure from brain activity by targeting the latent space of a pretrained image autoencoder, predicting semantic content by targeting learned vision-language embeddings, and combining both signals through a pretrained generative model that produces a final image reflecting both the recovered layout and the recovered meaning. Each notebook explains the reasoning behind its pipeline stage and provides runnable code to reproduce and build on each component. We present qualitative and quantitative metrics for all of our pipeline stages. Every notebook runs end-to-end on free-tier Google Colab hardware, and each stage can be inspected, modified, and replaced independently.

## 1 Introduction

When you look at a picture of a dog standing on a beach, your brain produces a structured pattern of activity. That pattern carries information about what you are seeing, from the spatial layout of the scene to the identity of the objects within it. A central question in visual neuroscience is how much of that information can be recovered from measured brain responses. Ultimately, can we reconstruct what a person was seeing from brain activity alone?

Early work established that visual content could, in fact, be read out from distributed fMRI activity patterns. Multi-voxel pattern analysis showed that responses in ventral temporal cortex contain enough information to discriminate viewed object categories^1,2^. Subsequent studies pushed beyond category-level decoding. Encoding models could identify which specific natural image a participant was viewing from a large candidate set^3^, and inverse retinotopic approaches could recover coarse spatial structure directly from activity in visual cortex^4,5^. Together, these studies showed that fMRI signals contain information sufficient to support not only discrimination and identification, but also constrained forms of visual reconstruction.

The introduction of deep neural networks substantially expanded what could be decoded and reconstructed. Rather than relying on hand-crafted stimulus features, researchers began using hierarchical visual representations extracted by trained networks as decoding targets^6,7^. In parallel, generative modeling approaches showed that learned image priors could improve reconstructions by supplying plausible visual detail that is not fully specified by the brain signal alone^8,9^. More recent work has also explored self-supervised objectives and related representation-learning strategies for reconstruction from brain activity^10^.

Over the past few years, many reconstruction pipelines have adopted a common high-level strategy that combines complementary types of decoded information. Low-level targets, such as autoencoding latents, preserve aspects of spatial layout, color, and coarse visual structure, but carry limited explicit information about object or scene identity. High-level targets, such as image embeddings derived from multimodal models like CLIP, capture abstract semantic content, but do not themselves constitute images^11,12^. Recent systems have shown that these signals can be combined within pretrained latent diffusion and related generative models to improve both structural fidelity and semantic accuracy^13–17^. This same general framework has also been extended toward shared-subject and cross-subject settings, reducing the amount of subject-specific data required for image reconstruction^18–21^.

Despite rapid progress, a substantial practical barrier remains. Many modern reconstruction pipelines are difficult to inspect, reproduce, or adapt. They often depend on large and complex codebases, require substantial computational resources, and involve multiple representational stages whose individual contributions are not always made explicit. As a result, a newcomer who reads a reconstruction paper and wants to understand how the system works, run it, and begin modifying it still faces a steep uphill climb.

The present work addresses this gap. We present a reproducible and quantitatively validated reference implementation for modern fMRI-to-image reconstruction in the Natural Scenes Dataset^22^. The pipeline is deliberately modular, with each stage isolating a recurring design decision in the reconstruction literature, including the choice of decoding target, the handling of repeated fMRI measurements, and the integration of decoded signals within a generative model. Each stage can therefore be understood, evaluated, and modified independently. Figure 1 summarizes the full workflow, and the remainder of the paper characterizes its methods, quantitative behavior, and empirical scope.

**Figure 1:**
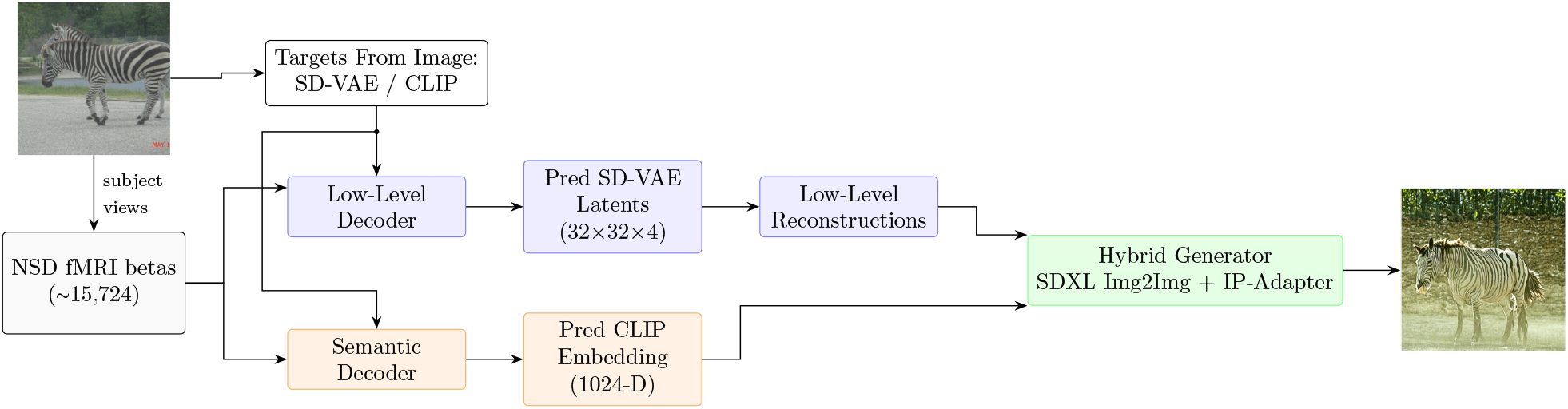
Overview of the reconstruction pipeline. Stimulus images are encoded into two complementary target spaces: SD-VAE latents (low-level) and CLIP embeddings (high-level). Separate decoders are trained to predict each target from fMRI responses. At generation time, the low-level reconstruction provides spatial structure, and the high-level embedding provides semantic guidance; both are combined through an SDXL image-to-image pipeline with IP-Adapter conditioning to produce the final output.

## 2 Methods

This work releases and evaluates a reference implementation for naturalistic stimulus reconstruction from fMRI in the Natural Scenes Dataset (NSD). The pipeline is modular, factoring the reconstruction problem into two decoded target spaces and a generative stage that combines them. Unless noted otherwise, the reported results correspond to a compute-constrained reference configuration using NSD subject 1 with all model settings chosen to fit within Google Colab free-tier hardware.

### 2.1 Pipeline Overview

As outlined, the pipeline factors reconstruction into three stages. The following sections specify each component.

#### Low-level target

The low-level path targets the latent space of the Stable Diffusion variational autoencoder (VAE)^12^. This encoder compresses each 256 × 256 RGB image into a 32 × 32 × 4 latent tensor containing 4,096 values, a roughly 48× reduction in dimensionality. The compression preserves global spatial layout, dominant colors, and coarse structural features while discarding fine texture and imperceptible detail. Because the latent representation is continuous and relatively low-dimensional, it is feasible to treat decoding as a standard regression problem at NSD scale, an approach also adopted by Ozcelik and VanRullen ^14^ using a similar autoencoder-based first stage. Figure 2 illustrates this encode-decode round trip on a single NSD stimulus.

**Figure 2:**
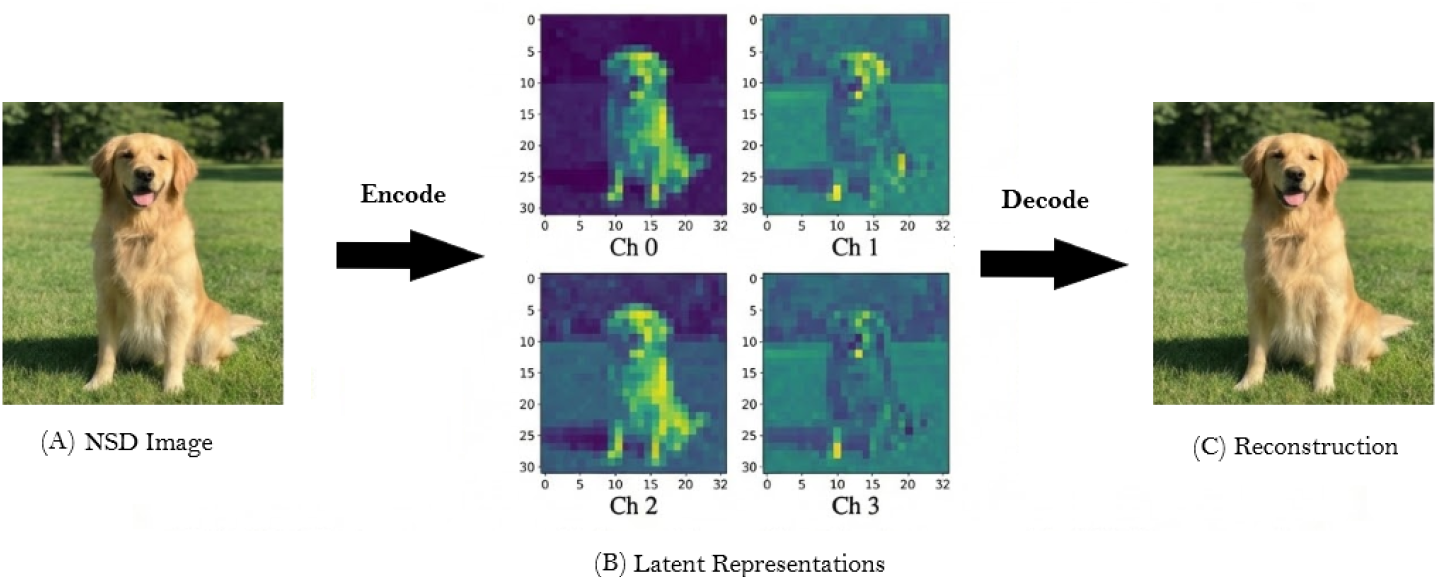
Illustration of the low-level target space. (a) An example NSD stimulus. (b) Its four-channel 32 × 32 latent tensor, visualized as separate heatmaps. (c) The image reconstructed by the VAE decoder. The near-lossless round trip confirms that the latent space preserves sufficient visual information to serve as a decoding target.

#### High-level target

The high-level path targets CLIP vision embeddings^11^, specifically from an OpenCLIP ViT-H/14 model trained on LAION-2B^23^. These embeddings compress each image into a 1,024-dimensional vector that emphasizes what is in the image (objects, scenes, categories) rather than exactly how it looks. Two photographs of different dogs in different settings will have similar CLIP embeddings because they depict the same concept, even though their pixel content is very different. CLIP embeddings cannot be directly turned back into pixels, but they provide a compact semantic code that can steer a generative model toward the right category of output. Figure 3 shows how this representation is extracted and evaluated.

**Figure 3:**
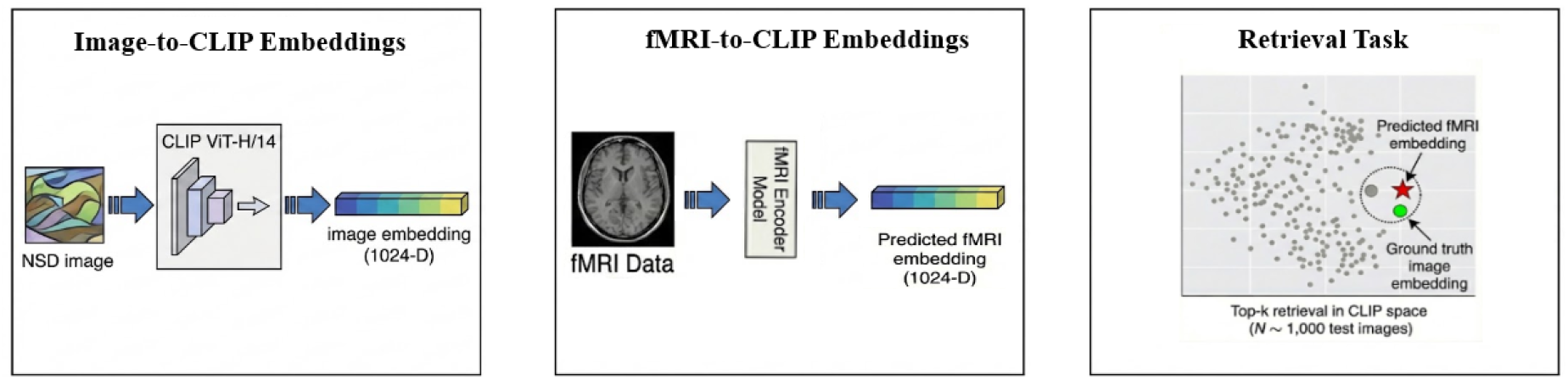
Overview of the semantic target and its evaluation. Each NSD stimulus image is encoded into a 1,024-dimensional CLIP embedding; the decoder predicts an embedding from fMRI; retrieval is evaluated by querying a pool of candidate image embeddings with the brain-predicted embedding.

#### Generative combiner

The final stage brings both signals together through an SDXL image-to-image pipeline. The low-level reconstruction is fed in as a starting image that tells the generative model where things are and roughly what colors to expect. The decoded CLIP embedding is injected through an IP-Adapter module^24^ that conditions the model’s cross-attention layers, telling it what objects and scenes should appear. Intuitively, the low-level signal provides the spatial skeleton and the high-level signal provides the semantic identity. This two-signal strategy follows the general approach used by recent end-to-end reconstruction systems^13–15^, while remaining compact enough for the chosen reference configuration.

### 2.2 Dataset and study design

All experiments use the Natural Scenes Dataset^22^, a large-scale 7T fMRI resource in which participants viewed tens of thousands of natural images drawn from COCO^25^ across dozens of scanning sessions. NSD provides pre-computed single-trial beta weights estimated from a general linear model, so the present work starts from preprocessed response estimates rather than raw BOLD timeseries. Each voxel value used in this work is already a single number summarizing the estimated response amplitude for one stimulus presentation.

The reference workflow uses the nsdgeneral mask, which restricts the roughly 200,000 whole-brain voxels to approximately 15,724 that are visually responsive, spanning early visual cortex through higher object-and scene-selective regions. This removes activity from regions that do not encode visual information and keeps the data small enough to fit in memory on modest hardware. Each image was shown to the subject three times in different scanning sessions, and practical choices such as how to handle repeated measurements, which decoder architecture and regularization to use, and how to balance generation parameters are explored in the corresponding modules. For the reported reference subject, the workflow uses 8,640 training images, 300 validation images, and 1,000 held-out test images. Figure 4 illustrates the voxel mask, an example activation pattern, and the data organization.

**Figure 4:**
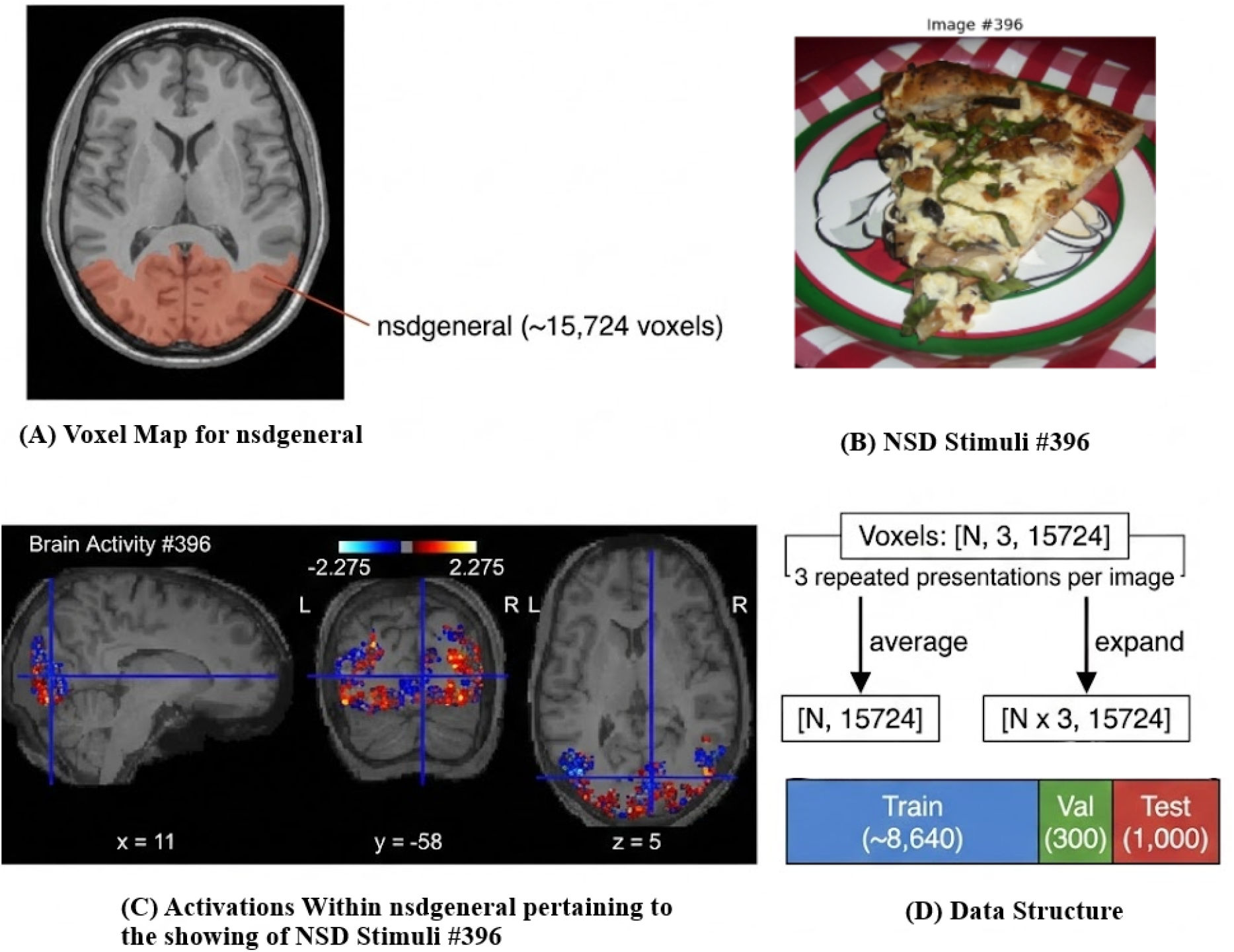
Practical NSD data setup. (A) The nsdgeneral voxel mask (about 15,724 voxels) overlaid on an anatomical slice. (B) Example NSD stimulus image. (C) Corresponding activation pattern within the nsdgeneral mask. (D) Tensor organization with three repeated presentations per image and the resulting train/validation/test splits.

### 2.3 Released implementation modules

The released resource is organized as six notebooks, including one setup notebook (Notebook 0) and five scientific modules that mirror the pipeline stages. Table 1 summarizes the notebooks and their outputs.

**Table 1:** Released implementation notebooks and primary outputs.

| Notebook | Stage | Primary output | Role in the pipeline |
| --- | --- | --- | --- |
| 0 | Colab primer and setup | Minimal Colab environment setup workflow | Provides a minimal GPU-enabled execution environment |
| 1 | fMRI and voxels | NSD loading and voxel-handling workflow | Defines the input representation and study split |
| 2 | VAE target preparation | Stable Diffusion VAE encode/decode workflow | Defines the low-level target space |
| 3 | Low-level decoding | Predicted VAE latents and decoded test images | Provides structural conditioning for generation |
| 4 | Semantic decoding | Predicted CLIP embeddings and retrieval metrics | Provides semantic conditioning for generation |
| 5 | Hybrid generation | Final reconstructions and ablation metrics | Combines both decoded signals |

The sequence is cumulative, and the first three modules are primarily conceptual. Notebook 0 provides a short Colab primer for readers who need a minimal execution environment. Module 1 asks what NSD fMRI data actually looks like in practice. It loads the beta weights for a single subject, applies the nsdgeneral voxel mask, organizes the repeated presentations into train, validation, and test splits, and saves the resulting arrays for use in all subsequent modules. By the end of this notebook, the reader has a concrete sense of the input data.

Module 2 explores the dimensionality mismatch between fMRI inputs and image outputs, and the problems that arise when trying to predict in one space from the other. It then introduces the Stable Diffusion VAE as a method for compressing images into a more learnable target space.

The remaining three modules each train a decoder, produce outputs, and evaluate them. The low-level decoder targets the latent space defined in Module 2, the semantic decoder targets CLIP embeddings, and the hybrid generator combines both decoded signals inside a generative model. The following sections describe each of these stages in detail.

### 2.4 Low-level decoder

The case against predicting pixels directly from brain activity is arithmetic. A 256 × 256 RGB image contains nearly 200,000 pixel values, but NSD provides only about 8,000 training images per subject and roughly 15,000 input voxels. That works out to approximately 0.04 training samples per output dimension, a regime in which overfitting is almost guaranteed and even small spatial misalignments between predicted and true images would be penalized heavily in pixel space, despite being perceptually insignificant.

The solution provided in this pipeline is to predict in a more compact representation. The Stable Diffusion VAE compresses each image into 4,096 continuous latent values, increasing the sample-to-dimension ratio to roughly two training examples per target dimension. While still not generous by machine learning standards, this is enough for regularized regression to produce meaningful results. Crucially, the latent representation preserves enough spatial and color information that decoded latents can be converted back to recognizable images through the VAE decoder. Moving the regression target into a space that is both learnable from fMRI and invertible back to image space is the key design choice at this stage.

Two decoder models are trained to make this prediction. The first is ridge regression, a linear model with L2 regularization that provides a stable baseline. Ridge is hard to overfit at NSD sample sizes and serves as a useful reference point, since anything the MLP achieves beyond ridge represents the value of modeling nonlinear relationships between voxels and latent features. The second model is a regularized multilayer perceptron (MLP), which can capture those nonlinear interactions but is also more sensitive to how the training data are presented. Predicted latents are decoded back to image space using the same VAE, and all evaluation is performed on 1,000 held-out test images that the decoders never saw during training. Because the decoder treats the VAE latent as a generic regression target, replacing Stable Diffusion’s autoencoder with a different one requires only changing the target extraction step in Notebook 2 while the decoder training in Notebook 3 remains the same.

### 2.5 Semantic decoder

The semantic decoder addresses a different question. Instead of recovering what the image looked like, can we recover what it was about? The target here is the CLIP embedding of the viewed image, a 1,024-dimensional vector that captures the semantic gist of the scene, including which objects, scenes, and categories are present.

As in the low-level stage, both a ridge regression baseline and an MLP are trained. Evaluation works differently here, though, because CLIP embeddings are not images and there is no natural way to look at the output and judge quality. Instead, the primary evaluation is framed as a retrieval task. The brain-predicted embedding is used as a search query against a pool of ground-truth image embeddings. Does the closest match correspond to the image the subject was actually viewing? Following Scotti et al. ^15^, success is measured by top-1 retrieval accuracy and pairwise accuracy over 300 held-out test images. As a complementary check, predicted embeddings are also passed through IP-Adapter to generate images conditioned solely on the decoded semantic signal, providing a qualitative sense of what information the embeddings carry before the hybrid stage combines them with low-level structure. The same pipeline structure applies to any fixed-dimensional image embedding, so swapping CLIP for a different representation such as DINOv2 requires changing only the feature extractor in Notebook 4.

### 2.6 Hybrid generation

The hybrid generator combines the two decoded outputs into a single image using an SDXL image-to-image pipeline implemented with the `diffusers` library. The low-level reconstruction, which is blurry but spatially informative, is upscaled to 1024 × 1024 using Lanczos resampling and provided as the starting image for the diffusion process. The decoded CLIP embedding is prepared for IP-Adapter conditioning by reshaping it into the format expected by the adapter and pairing it with a zero embedding for classifier-free guidance. We preserve the predicted embedding’s natural magnitude so that the conditioning more closely matches the scale seen by IP-Adapter during training.

Generation follows a two-pass strategy designed to balance semantic conditioning with structural preservation. The first pass uses stronger IP-Adapter conditioning and moderate diffusion strength to establish scene content while retaining the low-level layout. The second pass reduces the adapter scale and performs a lighter refinement pass on the first-pass output, improving detail without fully overriding the recovered structure.

All of this runs on a single T4 GPU with 16GB of VRAM, thanks to fp16 precision, VAE tiling to reduce peak memory during encoding and decoding, and a fast scheduler that achieves reasonable quality in a moderate number of denoising steps. All three conditions (low-level only, semantic only, and hybrid) are evaluated on 1,000 held-out test images using four metrics spanning pixel-level fidelity through high-level semantic similarity, following the evaluation protocol established by Scotti et al. ^15^. All generation parameters are exposed as arguments to the generation function in Notebook 5, allowing the reader to explore how diffusion strength, adapter scale, and guidance weight shift the balance between structural fidelity and semantic accuracy.

## 3 Results

### 3.1 Low-level reconstruction: recovering layout and color

How much spatial structure can a simple regression model recover from brain activity? Both the ridge baseline and the MLP (trained in Notebook 3) consistently recover the coarse spatial layout and dominant color palette of the original image, but fine-grained details like object boundaries, facial features, and texture are largely absent. The reconstructions typically look like heavily blurred, color-shifted approximations of the original.

Quantitatively, the MLP slightly outperformed ridge regression across the full 1,000-image test set, achieving a mean SSIM of 0.446 compared to 0.435 for ridge. To put these numbers in context, an SSIM of 1.0 would indicate a perfect pixel-level match, and a value around 0.4 reflects a reconstruction that preserves broad structure but differs substantially in detail. The MLP also produced higher SSIM on 800 of the 1,000 test images, confirming that the nonlinear model holds a consistent but modest advantage. PSNR and MSE followed the same pattern (MLP: 13.49 dB, Ridge: 13.29 dB), suggesting that the MLP’s improvement is not limited to structural similarity alone.

Figure 5 shows representative examples spanning a range of reconstruction quality, from cases where layout and color are clearly recognizable to cases where the output is largely uninformative. This variation is itself informative: images with strong color contrasts and simple compositions tend to decode well, while complex scenes with many small objects tend to produce diffuse, ambiguous reconstructions.

**Figure 5:**
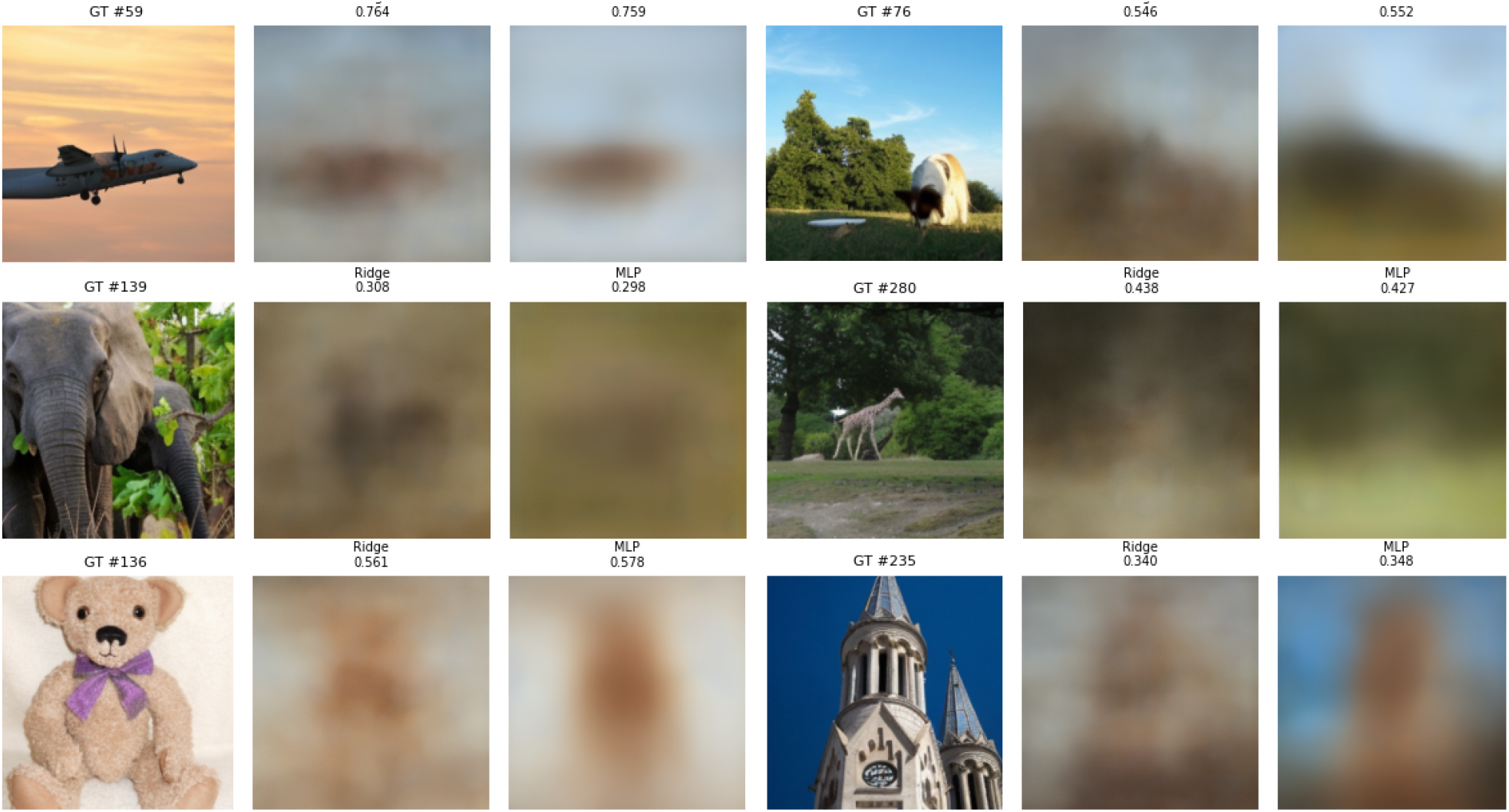
Low-level reconstruction results on the test set. For each stimulus, the figure shows the ground-truth NSD image together with ridge and MLP reconstructions obtained by predicting VAE latent representations from fMRI and decoding them back to image space. Numbers above the reconstructions report per-image SSIM. Both models often recover coarse color and global layout, while fine-grained structure and object identity remain weak.

**Figure 6:**
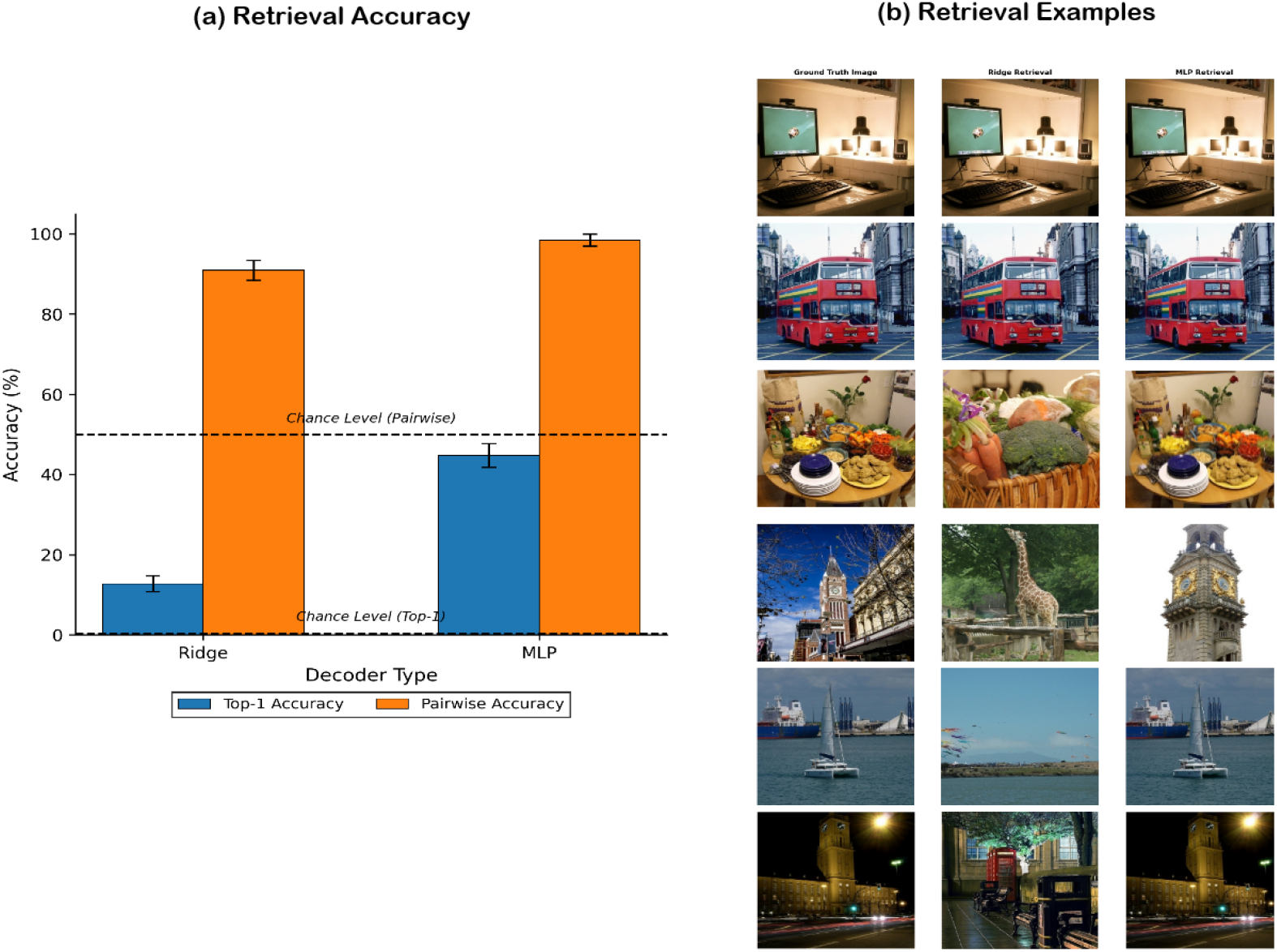
Semantic decoding performance. (a) Top-1 and pairwise retrieval accuracy on the first 300 test examples for ridge and MLP decoders, with chance level indicated. (b) Example retrievals showing the ground-truth image and the nearest neighbor in CLIP space based on the brain-predicted embedding.

### 3.2 Semantic decoding: recovering meaning

Where the low-level stage asks “what did the image look like?” the semantic stage asks “what was it about?”. The MLP decoder (trained in Notebook 4) achieved 45.67% top-1 retrieval accuracy on the first 300 test images, meaning that in nearly one out of two cases, the brain-predicted CLIP embedding was closer to the correct image embedding than to any of the other 299 candidates. Chance performance in this task is 1*/*300 ≈ 0.33%, so the decoder is well over 100 times above chance. The pairwise accuracy reached 98.63%, indicating that when given just two options, the brain-predicted embedding almost always picks the correct image over a random distractor.

Ridge regression also performed well above chance (17.0% top-1, 92.44% pairwise), but the MLP improved substantially over ridge. This is a much larger gap than in the low-level stage, where ridge and MLP were much closer together. The pattern suggests that the mapping from voxel patterns to semantic content is substantially nonlinear, and that even a relatively simple neural network can exploit this nonlinearity to meaningful effect.

An interesting qualitative pattern is that when the semantic decoder fails, it tends to fail gracefully: the retrieved image is often, although not always, semantically related to the correct one (e.g., retrieving a different outdoor scene rather than a completely unrelated image). This indicates that the decoded embeddings land in roughly the right neighborhood of CLIP space, even when they do not point to the exact correct image.

### 3.3 Hybrid reconstruction: combining structure and meaning

The hybrid stage (Notebook 5) tests whether combining low-level spatial structure with high-level semantic identity produces better reconstructions than either signal alone. We evaluate all three conditions across four metrics that capture different levels of visual similarity, following the protocol established by Scotti et al. ^15^. The results are summarized in Table 2.

**Table 2:** Hybrid reconstruction evaluation on 1,000 held-out test images at 425 × 425 resolution. PixCorr and SSIM measure pixel-level fidelity. InceptionV3 and CLIP measure semantic similarity as 2-way identification accuracy (chance = 50%). Higher is better for all metrics.

| Metric | Low-level only | Semantic only | Hybrid |
| --- | --- | --- | --- |
| PixCorr ↑ | .455 | .148 | .363 |
| SSIM ↑ | .520 | .362 | .371 |
| InceptionV3 ↑ | .519 | .954 | .941 |
| CLIP ↑ | .548 | .948 | .938 |

The four metrics span the range from raw pixels to abstract semantics. PixCorr and SSIM measure pixel-level fidelity, reflecting how closely the reconstruction matches the original in spatial structure and luminance patterns. InceptionV3 and CLIP operate at a higher level, measuring whether the reconstruction depicts the same kind of scene and objects as the original. These two neural network metrics are reported as 2-way identification accuracy. Given a reconstruction and two candidate images, one correct and one random distractor, how often does the model’s feature space pick the correct match? Chance performance is 50%. We note that pixel-level metrics here use 425 × 425 resolution following standard practice, while Section 3.1 reports at the native 256 × 256.

The pattern across conditions is consistent. Low-level-only reconstructions dominate on pixel-level metrics because they directly preserve spatial structure from the decoded latents, but they remain weak on high-level metrics because a blurry color-matched image often lacks clear object identity. Semantic-only generation flips this: high-level scores are very strong, but pixel-level fidelity collapses because the generative model has no spatial anchor and invents layouts freely. The hybrid pipeline occupies the middle ground on every metric. It gives up some of the spatial fidelity of the low-level path, but retains much more structure than the semantic-only condition while approaching semantic-only performance on both InceptionV3 and CLIP. No single condition dominates across all four metrics, which is precisely the tradeoff the hybrid approach is designed to navigate.

Qualitative examples in Figure 7 make this visible. The low-level reconstructions are blurry and category-ambiguous: a blob of brown and green might be a dog, a field, or a piece of furniture. When semantic conditioning is added, the generative model resolves this ambiguity, producing an output that is recognizably a dog, a building, or a person, while the low-level input prevents the model from inventing a spatial layout that contradicts the brain signal.

**Figure 7:**
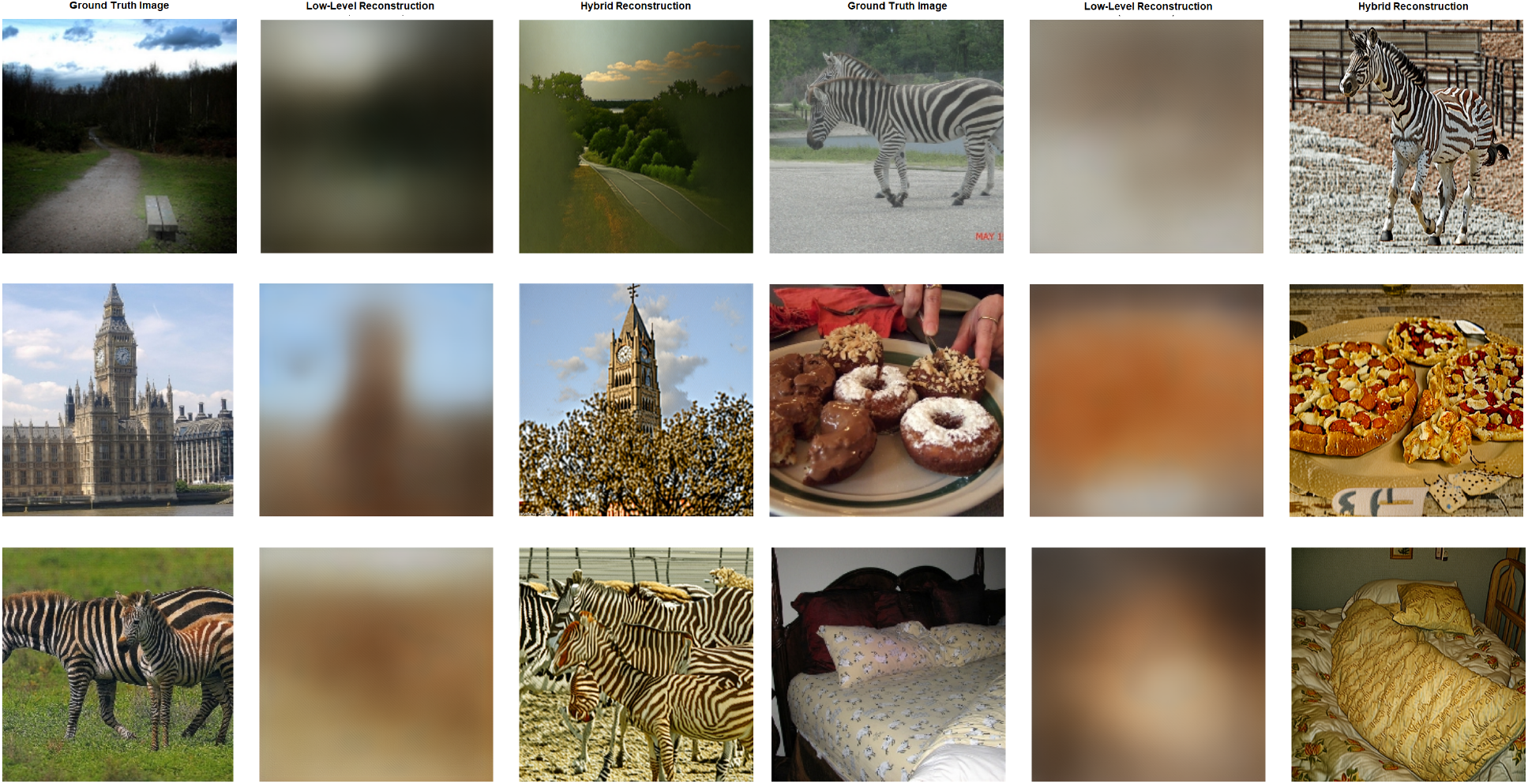
Hybrid reconstructions on held-out test stimuli. Columns 1 and 4 show the ground truth, 2 and 5 show the low-level reconstruction, and 3 and 6 show the hybrid output that combines low-level structure with CLIP-based semantic conditioning.

## 4 Discussion

The primary contribution of this work is pedagogical, not architectural. The pipeline is designed to make each stage of modern fMRI-to-image reconstruction transparent, modifiable, and runnable on accessible hardware. Systems such as MindEye2^19^ and Brain-IT^26^ achieve higher overall reconstruction quality through shared-subject training, more expressive decoder architectures, and considerably more compute, but the present pipeline nonetheless produces competitive results across both pixel-level and semantic metrics.

### 4.1 Positioning relative to published systems

To give readers a sense of where the present pipeline falls relative to the field, Table 3 places our hybrid results alongside published numbers from recent reconstruction systems, all evaluated on NSD Subject 1 at 425 × 425 resolution following the standard protocol.

**Table 3:** Comparison with published reconstruction systems. All methods use NSD Subject 1 evaluated at 425 × 425.^†^.

| Method | PixCorr | SSIM | Incep. | CLIP |
| --- | --- | --- | --- | --- |
| Takagi & Nishimoto<sup>a</sup> | - | - | .760 | .770 |
| Brain-Diffuser<sup>b</sup> | .305 | .367 | .878 | .925 |
| MindEye<sup>c</sup> | .390 | .337 | .945 | .946 |
| MindEye2<sup>d</sup> | .374 | .439 | .962 | .934 |
| Ours | .363 | .371 | .941 | .938 |

<sup>a</sup>Takagi and Nishimoto<sup>13</sup>
<sup>b</sup>Ozcelik and VanRullen<sup>14</sup>
<sup>c</sup>Scotti et al.<sup>15</sup>
<sup>d</sup>Scotti et al.<sup>19</sup>
<sup>†</sup>Some recent systems are omitted because they report only multi-subject averages rather than per-subject results.

The results show that our pipeline is competitive with published systems across both pixel-level and semantic metrics despite using simpler components. In particular, the final hybrid system achieves strong semantic alignment while remaining reasonably faithful to the decoded low-level structure. We also note that generation-stage hyperparameters such as diffusion strength, guidance scale, and IP-Adapter conditioning weight can shift the balance between pixel-level and semantic scores considerably, and even small changes in these values can produce noticeably different metric profiles.

### 4.2 Scope and limitations

Several limits follow directly from the decision to present a compact reference configuration that fits within free-tier compute. The workflow starts from preprocessed NSD beta estimates and does not address upstream fMRI preprocessing. All results are reported for a single subject rather than a shared-subject model. Decoder architectures are intentionally simple and optimized for legibility and fast training rather than peak performance. These choices favor accessibility and reproducibility, and researchers with access to greater compute can extend the pipeline without modifying the core workflow since the generation loop is parametric over the number of test samples.

Beyond these pipeline-specific choices, reconstruction approaches built on NSD face broader challenges. The dataset was collected at 7 Tesla with 1.8 mm resolution across dozens of sessions per subject, providing signal quality that most scanning environments cannot match. Any reconstruction pipeline that relies on a pretrained generative model also inherits that model’s learned priors about what natural images look like, and disentangling what the decoder actually recovered from brain activity from what the generative model filled in remains an open challenge across the field. More fundamentally, whether these methods generalize to other image domains, scanning protocols, or subject populations is not yet well understood.

### 4.3 Practical value and extensibility

Despite these limits, the released workflow is designed for modification. Because the low-level, semantic, and hybrid stages are cleanly separated, each stage can be replaced without rebuilding the surrounding infrastructure. The Colab-first execution path also makes the resource suitable for teaching, reproduction exercises, and baseline development in settings where dedicated hardware is not available.

## 5 Conclusion

This work provides a reproducible, end-to-end primer for naturalistic stimulus reconstruction from NSD fMRI data. By factoring the problem into low-level latent decoding, high-level semantic decoding, and generative hybrid combination, the pipeline makes explicit how modern reconstruction systems are built from interpretable, composable parts. The low-level path recovers spatial layout and color, the high-level path recovers object and scene identity, and the generative combiner synthesizes both into a coherent image that is richer than either signal alone. The released implementation is designed first to teach and second to serve as a starting point, giving researchers a working system they can inspect, modify, and build on as the field continues to advance.

## Ethics statement

This study analyzed publicly available, de-identified data from the Natural Scenes Dataset. Original NSD data collection was conducted with informed consent from participants and under the ethics procedures reported in Allen et al. ^22^. No new human data were collected for the present study.

## Data and Code Availability

The Natural Scenes Dataset is publicly available at http://naturalscenesdataset.org^22^. Released notebooks and associated utility code are available at https://github.com/UYildiz12/nsd-tutorial-notebooks and https://drive.google.com/drive/folders/1cfp56-PGr29HwVtIa1DPWKzSPFoxBDcp?usp=sharing. A short Colab primer (Notebook 0) is included for readers who need a minimal GPU-enabled execution environment but is not part of the scientific content of the paper.

All notebooks have been tested end-to-end on Google Colab free-tier hardware (T4 GPU) as of February 2026. Approximate runtimes are summarized in Table 4; actual times vary with network speed during data streaming and model downloads.

**Table 4:**
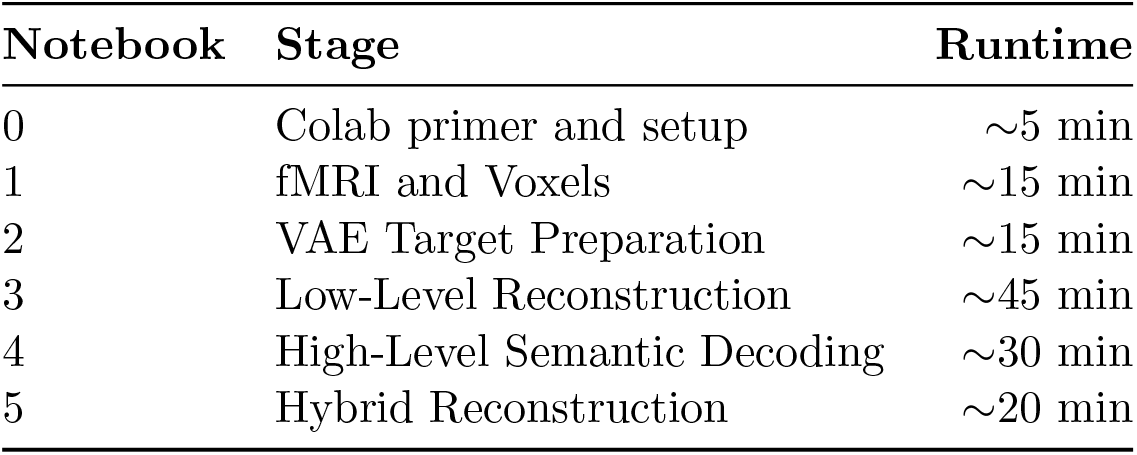
Approximate runtimes on Google Colab (T4 GPU).

## CRediT author statement

Umur Yildiz: Conceptualization, Methodology, Software, Formal analysis, Investigation, Visualization, Writing – original draft. Burcu A. Urgen: Conceptualization, Methodology, Supervision, Writing – review & editing.

## Acknowledgements

The authors thank the Natural Scenes Dataset team for making the dataset publicly available.

